# A novel germline mutation in the *POT1* gene predisposes to familial non-medullary thyroid cancer

**DOI:** 10.1101/2020.03.23.004663

**Authors:** Aayushi Srivastava, Beiping Miao, Diamanto Skopelitou, Varun Kumar, Abhishek Kumar, Nagarajan Paramasivam, Elena Bonora, Kari Hemminki, Asta Försti, Obul Reddy Bandapalli

## Abstract

Non-medullary thyroid cancer (NMTC) is a common endocrine malignancy with a genetic basis that has yet to be unequivocally established. In a recent whole genome sequencing study of five families with recurrence of NMTCs, we shortlisted promising variants with the help of bioinformatics tools. Here, we report *in silico* analyses and *in vitro* experiments on a novel germline variant (p.V29L) in the highly conserved oligonucleotide/oligosaccharide binding domain of the *Protection of Telomeres 1 (POT1)* gene in one of the families. The results showed that the variant demonstrates a reduction in telomere-bound POT1 levels in the mutant protein as compared to its wild-type counterpart. HEK293Tcells carrying *POT1*^*V29L*^ showed increased telomere length in comparison to wild type cells, strongly suggesting that the mutation causes telomere dysfunction and may play a role in predisposition to NMTC in this family. This study reports the first germline *POT1* mutation in a family with a predominance of thyroid cancer, thereby expanding the spectrum of cancers associated with mutations in the shelterin complex.

## Introduction

Thyroid cancer is the most frequently diagnosed malignant endocrine tumor with a world average age-standardized incidence rate of 6.7/100,000 persons per year (Bray, Ferlay et al., 2018). Non-medullary thyroid carcinoma (NMTC) accounts for up to 95% of all thyroid cancers (Hincza, Kowalik et al., 2019, Peiling Yang & Ngeow, 2016). Based on the population-based registers of the Nordic countries the risk of NMTC is about three-fold higher when a first-degree relative is diagnosed with NMTC compared to those without affected family members (Fallah, Pukkala et al., 2013). Apart from the rare syndromic forms of familial NMTC (FNMTC), including familial adenomatous polyposis, Gardner syndrome, Cowden syndrome, Carney complex type 1, Werner syndrome, and *DICER1* syndrome, the genetic basis of FNMTC is largely unknown (Hincza et al., 2019, Peiling Yang & Ngeow, 2016). FNMTC has been associated with an earlier age of onset, a higher incidence of multifocality and more aggressive disease compared to its sporadic counterpart (El Lakis, Giannakou et al., 2019, Fallah et al., 2013). Thus, it is important to identify genetic factors behind the familial disease to facilitate genetic counseling and clinical management of the patients.

Various approaches, including genome-wide association studies, linkage analyses, targeted sequencing, and whole exome sequencing, have been employed to gain understanding into the genetic basis of FNMTC. Several genes and loci, including mainly low-penetrance variants near or in *FOXE1, SRGAP1, TITF-1/NKX2-1, DIRC3, and CHEK2*, have been suggested to affect non-syndromic FNMTC susceptibility (Hincza et al., 2019, Peiling Yang & Ngeow, 2016). In addition, an imbalance of the telomere-telomerase complex has been demonstrated in the peripheral blood of familial papillary thyroid cancer patients (Capezzone, Cantara et al., 2008).

Recently, we performed whole genome sequencing (WGS) on five families with documented recurrence of NMTC and analyzed these samples with our in-house developed variant prioritization pipeline (FCVPPv2) along with other *in silico* tools (Srivastava, Kumar et al., 2019). This allowed us to identify a novel missense variant (p.V29L) in the *protection of telomeres 1* (*POT1*) gene in one of the families. POT1 is a critical component of the shelterin complex, which binds and protects telomeres by modulating telomere capping, replication, and extension by telomerase (de Lange, 2018). Structurally, it is the only member of the shelterin complex that contains two N-terminal oligonucleotide/oligosaccharide binding (OB) domains, which can bind the single-stranded TTAGGG repeats as well as a C terminus, which can bind to TPP1, anchoring it to the shelterin complex composed of four other components: TRF1, TRF2, TIN2 and RAP1 (de Lange, 2018). Germline variants in *POT1* have been described in familial melanoma (Potrony, Puig-Butille et al., 2019, Robles-Espinoza, Harland et al., 2014, Shi, Yang et al., 2014, Wong, Robles-Espinoza et al., 2019), glioma (Bainbridge, Armstrong et al., 2015), Li-Fraumeni-like syndrome (Calvete, Martinez et al., 2015b), colorectal cancer (Chubb, Broderick et al., 2016), chronic lymphocytic leukemia (Speedy, Kinnersley et al., 2016) and Hodgkin lymphoma (McMaster, Sun et al., 2018).

In this study, we elucidated the genetic and functional consequences of the novel *POT1* missense variant p.V29L that segregated in an NMTC family. Results from *in silico* and *in vitro* analyses suggested a mechanism for the identified *POT1* mutations in predisposition to FMNTC.

## Results

### WGS and variant prioritization

The subject of this study was an Italian family with reported recurrence of NMTC (Fig. 1). Five members of this family were affected by PTC, Hürthle cell cancer, micro-PTC or a combination of two subtypes (II-2, II-3, II-5, II-8, II-9). Three members were possible carriers affected by benign nodules (I-1, II-4, II-6) and two were unaffected (II-1, II-7). WGS was performed on eight of these family members. The variants were filtered based on pedigree data considering family members diagnosed with NMTC or micro-PTC as cases, benign nodules or goiter as potential variant carriers and unaffected members as controls.

**Figure 1A.**
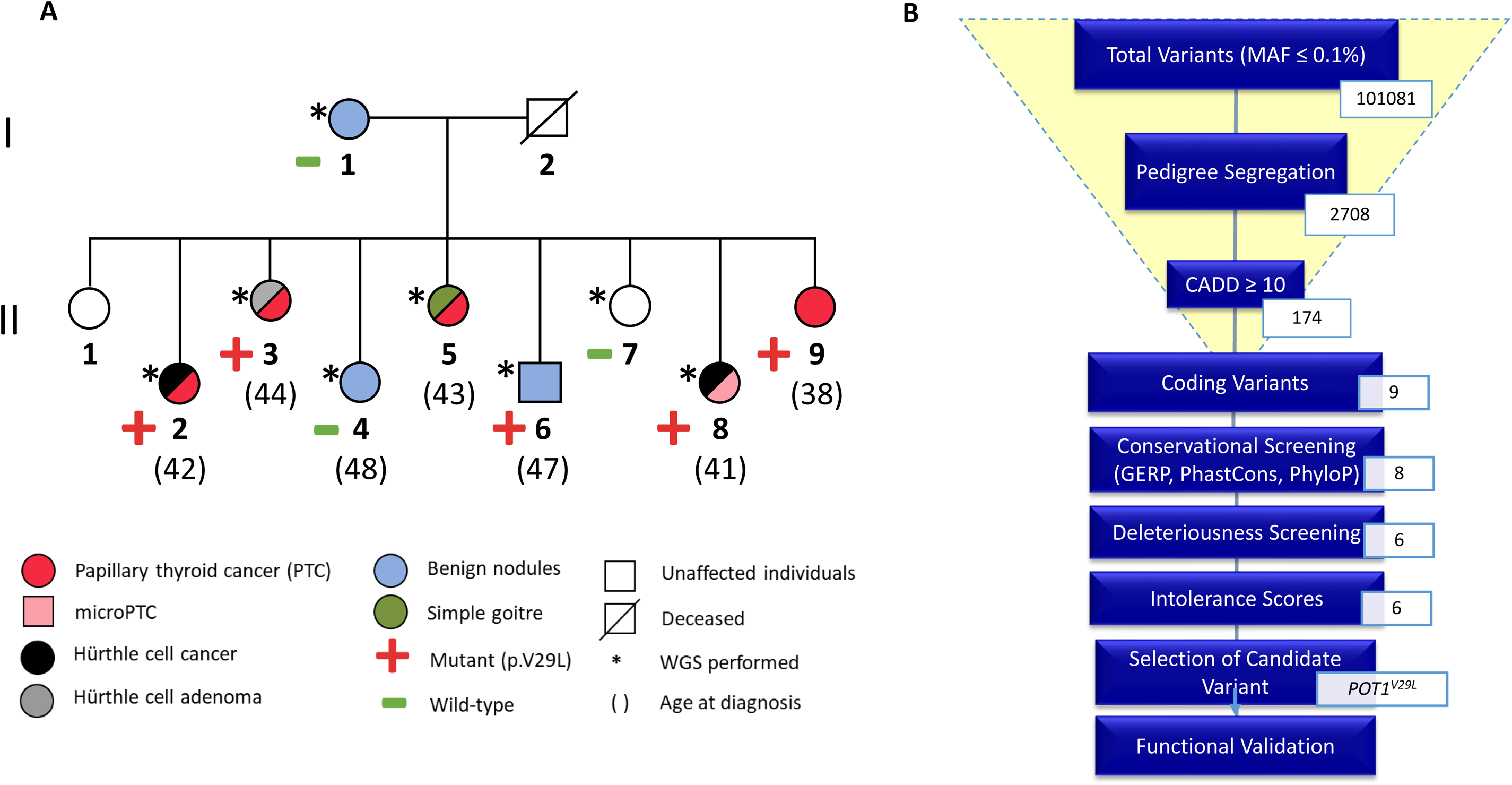
Pedigree of the NMTC family with *POT1*^*V29L*^ mutations. **Figure 1B.** Overview of the variant filtering process using the Familial Cancer Variant Prioritization Pipeline version 2 (FCVPPv2). The number of variants passing each step of the pipeline is shown.

A total of 101081 variants, with mean allele frequencies less than 0.1%, was reduced by pedigree-based filtering to 2708. We did not identify any deleterious loss-of-function variants, however, six non-synonymous variants in six genes (*EPYC, SPOCK1, MYBPC1, ACSS3, NRP1, and POT1*) segregated with the disease in the family and passed the filters of the FCVPPv2 (Srivastava et al., 2019). An overview of the process leading to the selection of a candidate variant is outlined in Figure 1B. Given the importance of *POT1* in various cancers, we selected it as our candidate variant for further *in silico* analyses and functional validation. A list of all shortlisted variants and their scores is available in the supplementary data (Table S1).

### *In silico* studies predict the importance of the p.V29L mutation to POT1 protein function

Comparative sequence analysis of the p.V29L position showed it to be highly conserved across selected representative species within the phylogeny (Fig. 2A). The p.V29L variant is located in the OB1 domain of the protein, as are several other germline and somatic variants reported in a wide spectrum of other human cancers (Fig. 2B, Table S2). It is also evident that the region around the position p.V29L is highly conserved. The tolerance of POT1 protein function to single amino acid substitutions was calculated by SNAP2 and accessed using PredictProtein. The heat map representation of the resulting data shows a highly deleterious effect of almost all substitutions in the position p.V29. An aggregation of highly deleterious effects of any amino acid change can be seen in the selected range (1-72 amino acids; Fig. 2C). These predictions reinforce the biological importance of the OB folds.

**Figure 2.**
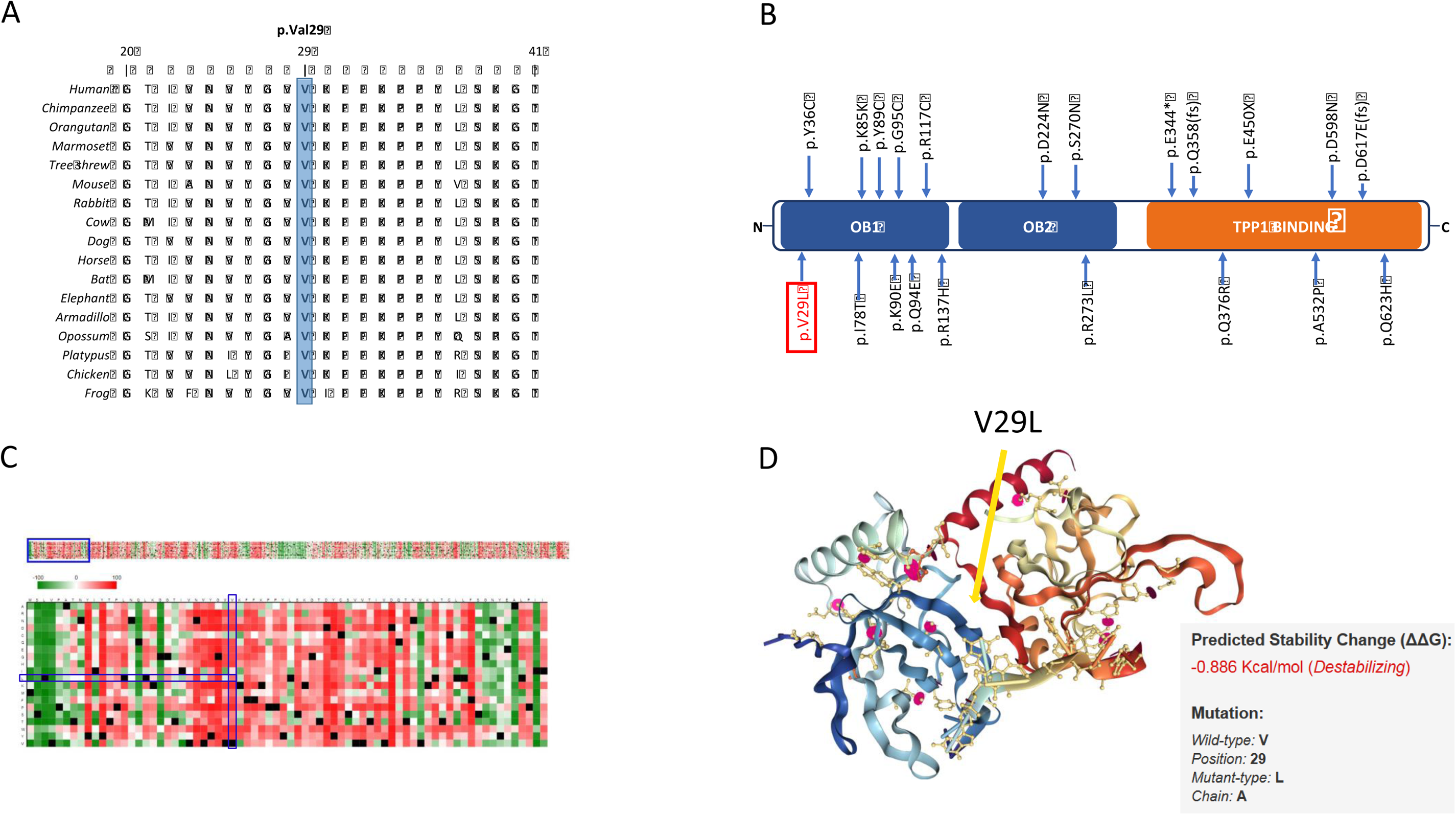
*In silico* studies. (A). Comparative sequence analysis of *POT1* across representative phylogeny. (B). Schematic primary structure of the POT1 protein with known germline mutations identified in various cancers shown relative to the OB domains (blue) and the TPP1 binding region (orange). The variant identified in this study is highlighted in red. (C). Heat map representation of SNAP2 results showing the predicted impact of individual amino acid substitutions (y-axis) for each position (x-axis) on protein function. Dark red indicates a highly deleterious substitution (score = 100), white indicates a minor effect, green indicates a neutral effect or no effect (score = -100), black represents the corresponding wild type residue (upper panel). The section of amino acids belonging to the OB1 domain from the upper panel is expanded and displayed in detail in the lower panel. p.V29L is shown with the blue rectangles (lower panel). (D). Crystal structure of the N-terminal domain of POT1 as a complex with ssDNA (PDB 1XJV). V29L is indicated with an arrow. Stability change caused by the p.V29L substitution as predicted by mCSM.

We attained the crystal structure of the N-terminal domain of POT1 (aa 1-185) as a complex with ssDNA from the RCSB PDB database (1XJV) (Lei, Podell et al., 2003). This domain binds G-rich telomeric ssDNA with the same specificity and higher affinity than the full-length protein, suggesting that this segment encompasses the entire DNA-binding region of the protein (Lei et al., 2003). The OB-fold shown in this crystal structure consists of a highly curved, five-stranded anti-parallel β-barrel. The interaction of the ssDNA with the concave groove of the OB folds along with the position of our variant (p.V29L) can be seen in Figure 2D. Moreover, we predicted the change in protein stability by the p.V29L substitution using the mutation Cutoff Scanning Matrix approach (mCSM), which relies on graph-based signatures to predict the impact of missense mutations on protein stability. The thermodynamic change in free energy caused by the p.V29L mutation was predicted to be destabilizing (ΔΔG = -0.886 Kcal/mol) (Fig. 2D).

### The V29L mutation aggravates DNA dependent functions of POT1

As adverted to in the introduction, both germline and somatic deleterious mutations reported in *POT1* tend to be concentrated in its OB folds (Gu, Wang et al., 2017). Therefore, these OB domains are the main target of the mutational events in this protein across different human cancers. Mutations in this region have been reported to affect DNA binding and lead to loss of function of the POT1 protein (Calvete, Martinez et al., 2015a, Ramsay, Quesada et al., 2013). To test whether the missense *POT1* variant affects protein function, we performed western blotting with lysates isolated from HEK293T cells transfected with an empty vector, or with the vector carrying cDNA encoding Myc-tagged human wild-type POT1 or Myc-tagged human mutant POT1. We did not detect significant differences in POT1 protein levels between POT1^WT^ and POT1^V29L^ transfected cells (Fig. 3A). We then performed chromatin immunoprecipitation (ChIP) assays to examine the effect of the POT1^V29L^ variant on the binding of POT1 to telomeric chromatin. Our results showed significantly weakened binding of telomeric DNA to POT1^V29L^ as compared to POT1^WT^ (Fig. 3B).

**Figure 3.**
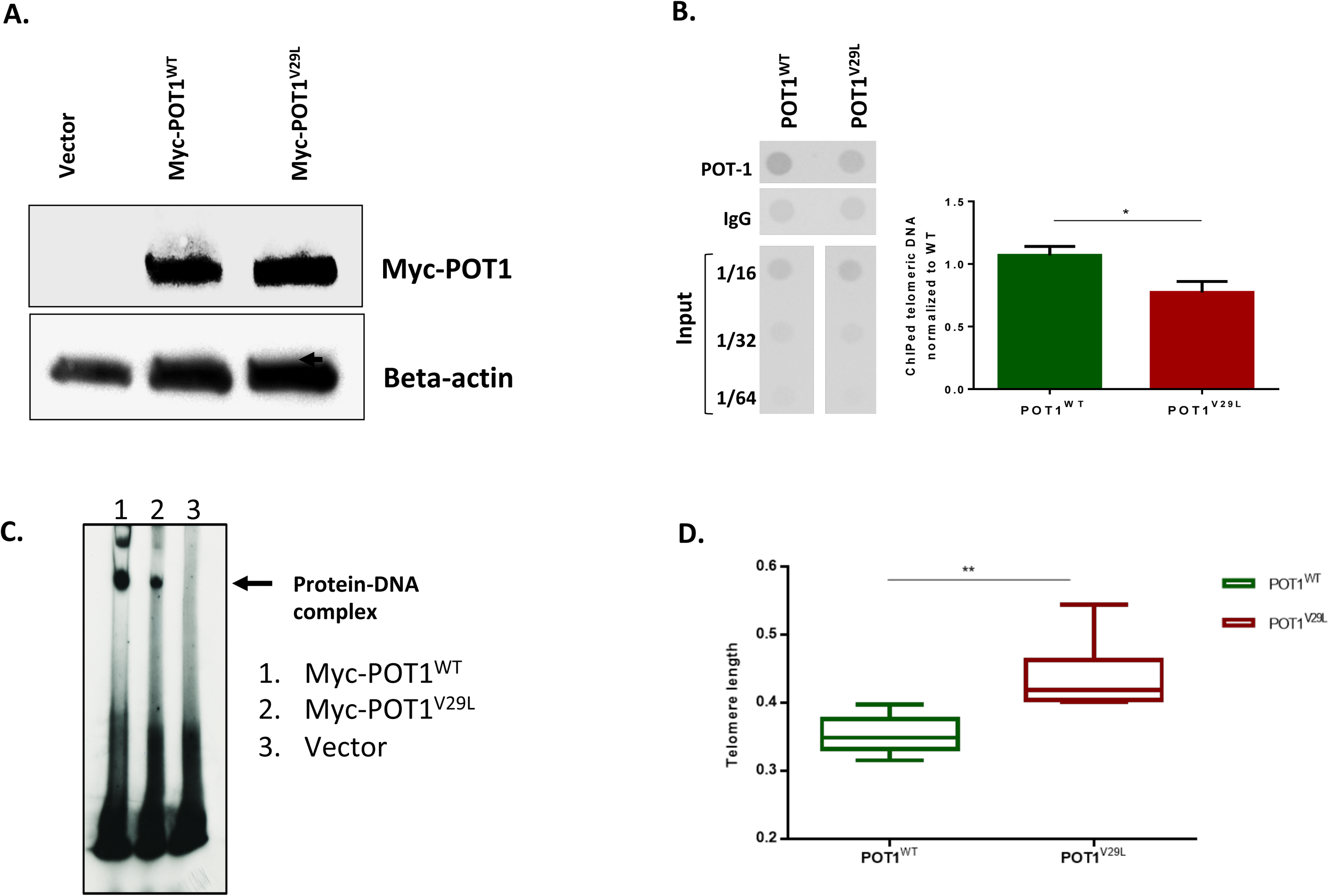
The *POT1*^*V29L*^ mutation affects telomeric binding to ssDNA. (A) Western blot of empty (untagged) vector, Myc-POT1 WT and Myc-POT1 mutant with beta-actin serving as an internal control. (B) Quantification of telomeric DNA bound to POT1 by ChIP analysis. IgG served as a negative control. Results were normalized to input chromatin. Representative ChIP dot blot is shown with input sample dilutions (left panel). Quantification of ChIP resulted in the bar graph (right panel). Green bar: wild-type; red bar: mutant. (C) Results of EMSA showing the decrease in POT1 binding capacity to telomeric ssDNA by the p.V29L substitution. (D) Relative telomere length is significantly longer in POTV29L compared to POT1WT transfected HEK293T cells after 40 passages.

Furthermore, to confirm our findings from the ChIP assay, we performed an electrophoretic mobility shift assay (EMSA) using constructs containing cDNA for wild-type and mutant POT1 that were translated *in vitro* and incubated with radiolabeled telomeric ssDNA. EMSA results confirmed that the p.V29L alteration affected the ability of POT1 to bind to the 3’ end of the G-rich telomeric overhang, whereas wild-type POT1 was able to efficiently bind to telomeric ssDNA (Fig. 3C). In an attempt to assess the effect of the POT1^V29L^ variant on telomere length, we measured telomere length in the family members using WGS data and found no significant difference. This could be due to naturally occurring variance in telomere length within the human population as well as due to age differences and the limited number of samples. This situation is distinct from one that is present in the analysis of cell lines. We demonstrated this experimentally by passaging 40 times of HEK293T cells transfected with POT1^WT^ and POT1^V29L^ and subsequently re-analyzing the telomere lengths. The results showed that the telomere length was significantly longer in mutated cells compared to the wild-type cells (two-tailed student’s t-test, P<0.005; Fig. 3D).

## Discussion

In our study, we identified a novel germline *POT1* missense mutation that segregated with thyroid cancer in an Italian family. As the scope of personalized therapy and medical genetics advance, the importance of identifying mutations and pathways affected in different cancers is heightened. Next-generation sequencing has emerged as the state-of-the-art tool for the identification of driver mutations in tumors and novel cancer-predisposing genes in Mendelian diseases. The heritability of thyroid cancer can be attributed to both rare, high-penetrance mutations and common, low-penetrance variants. Our approach was focused on identifying the former in a familial pedigree.

The *POT1* variant prioritized in this family (p.V29L) underwent several *in silico* and *in vitro* studies to demonstrate the consequence of the amino acid substitution. Since the mutation is located in the OB1-domain, a thorough literature review aided us in hypothesizing the functional effects of the point mutation (Gu et al., 2017), as several other germline and somatic variants reported in a wide spectrum of other human cancers are also clustered around the OB-domains. These predictions were supported by results from both the *in silico* studies as well as the *in vitro* studies. *In silico* studies showed putative disruption of POT1 protein function by p.V29L that was later validated by functional studies. The ChIP assay showed a significant decrease in the mutant POT1 protein’s ability to bind to ssDNA as compared to its wild-type counterpart (p= 0.01, student’s t-test.). Results from the EMSA supported this inference by also showing a decreased ability of *POT1*^*V29L*^ in forming a protein-DNA complex. Although we did not test the capability of the *POT1*^*V29L*^ protein to interact and bind to TPP1, thus allowing it to localize to double-stranded telomeres, a previous functional study on a variant in the OB-folds of POT1 (p.R117C) showed disruption in POT1-TPP1 interaction as a result of the mutation (Calvete, Garcia-Pavia et al., 2017).

Germline deleterious mutations in *POT1* have previously been associated with susceptibility to melanoma (Robles-Espinoza et al., 2014, Shi et al., 2014), glioma (Bainbridge et al., 2015), colorectal cancer (Chubb et al., 2016), Li-Fraumeni-like syndrome (Calvete et al., 2015b) and chronic lymphocytic leukemia (Speedy et al., 2016). Orois et al. analyzed seven FNMTC families with the sole aim of identifying *POT1* mutations but were unable to detect any variants in these families (Orois, Badenas et al., 2020). It is of particular interest to note that predicting the phenotype simply by attaining the genotype is not possible. Mutations in the same domain of the POT1 protein can lead to an array of different human cancers.

Although advancements have been made in recent years in the understanding of FNMTC, the hereditary factors contributing to the susceptibility to and possible unfavorable prognosis of FMNTC have yet to be adequately explored. On the one hand, over-diagnosis and overtreatment of low-grade disease or benign nodules have to be avoided, and on the other hand, it is imperative to identify aggressive cases with poor prognoses (Jegerlehner, Bulliard et al., 2017). This is only possible if there is a strong understanding of predictive germline variants and their underlying pathways. We acknowledge that one of the limitations of this study is that the proposed disease-causing variant was found in only one family. However, when dealing with rare, high-penetrance variants, it is a challenging task to locate more than one family with a mutation in the same gene. Nonetheless, this draws attention to two aspects. First, it is evident that many other disease-causing loci have yet to be discovered and second, there is a certain ambiguity associated with the selection of one causal variant in a family, as other deleterious variants that are shared amongst patients in the family could also be important in the pathogenesis of the studied phenotype.

In conclusion, the *POT1* mutation reported in this study plays a role in NMTC predisposition. While one germline mutation in *POT1* has already been reported in a melanoma-prone family with recurrence of thyroid cancers (Wilson, Hattangady et al., 2017), we report the first of such mutations in a family affected solely by NMTCs. Hence, our study expands the spectrum of cancers known to be evoked by inherited mutations in the *POT1* gene. Loss of function of this gene may play a role in the pathogenesis of NMTC and may facilitate novel approaches for screening and clinical management of this disease.

## Materials and Methods

### Patients

The subject of this study was an Italian family with NMTC aggregation recruited at the S. Orsola-Malpighi Hospital, Unit of Medical Genetics in Bologna, Italy. Samples from four affected members, one unaffected member, and three possible carriers were available for whole genome sequencing. All blood samples were collected from the participants with informed consent following ethical guidelines approved by “Comitato Etico Indipendente dell ‘Azienda Ospedaliero-Universitaria di Bologna, Policlinico S. Orsola-Malpighi (Bologna, Italy)” and “comité consultatif de protection des personnes dans la recherche biomédicale, Le centre de lutte contre le cancer Léon-Bérard (Lyon, France)” and DNA was isolated using the QiAMP DNA Blood Mini kit according to the manufacturer’s instructions.

### Whole Genome Sequencing

WGS of available DNA samples from the NMTC family members was performed using Illumina-based small read sequencing. Mapping to the human reference genome (assembly GRCh37 version hs37d5) was performed using BWA mem (version 0.7.8) (Li & Durbin, 2009) and duplicates were removed using biobambam (version 0.0.148). Platypus (Rimmer, Phan et al., 2014) was used to call small nucleotide variants (SNVs) and InDels through joint calling on all the samples from the family. Variants were annotated using ANNOVAR, 1000 Genomes, dbSNP and ExAC (Lek, Karczewski et al., 2016, Smigielski, Sirotkin et al., 2000, The Genomes Project (Auton et al., 2015, Wang, Li et al., 2010). Variants with a QUAL score greater than 20 and coverage greater than 5x, and that passed all the Platypus internal filters were evaluated further. Variants with minor allele frequencies (MAFs) greater than 0.1% in the 1000 Genomes Phase 3 and non-TCGA ExAC data were marked as common and removed. A pairwise comparison of shared rare variants was performed to check for sample swaps and family relatedness.

### Data Analysis and Variant Prioritization

#### Prioritization of variants

Variant evaluation was performed using the criteria of our in-house developed variant prioritization pipeline FCVPPv2 (Kumar, Bandapalli et al., 2018). First, all the variants were filtered based on the pedigree data considering cancer patients as cases, individuals with benign nodules as potential mutation carriers and unaffected persons as controls.

Variants were then filtered using the Combined Annotation Dependent Depletion (CADD) tool v1.3 (Kircher, Witten et al., 2014), and variants with a scaled PHRED-like CADD score greater than 10, i.e. variants belonging to the top 10 % of probable deleterious variants in the human genome, were considered further. Genomic Evolutionary Rate Profiling (GERP) (Cooper, Stone et al., 2005), PhastCons (Siepel, Bejerano et al., 2005) and PhyloP (Pollard, Hubisz et al., 2010) were used to evaluate the evolutionary conservation of the genomic position of a particular variant. GERP scores > 2.0, PhastCons scores > 0.3 and PhyloP scores ≥ 0.3 were indicative of a good level of conservation and were therefore used as thresholds in the selection of potentially causative variants.

Next, all variants were assessed for deleteriousness using 10 tools accessed using dbNSFP (Liu, Wu et al., 2016), namely SIFT, PolyPhen V2-HDV, PolyPhen V2-HVAR, LRT, MutationTaster, Mutation Assessor, FATHMM, MetaSVM, MetLR and PROVEAN and variants predicted to be deleterious by at least 60% of these tools were analyzed further.

Lastly, three different intolerance scores derived from NHLBI-ESP6500 (Petrovski, Wang et al., 2013), ExAC (Lek, Karczewski et al., 2016) and a local dataset, all of which were developed with allele frequency data, were included to evaluate the intolerance of genes to functional mutations. The ExAC consortium has developed two additional scoring systems using large-scale exome sequencing data including intolerance scores (pLI) for loss-of-function variants and Z-scores for missense and synonymous variants. These were used for nonsense and missense variants respectively. However, all the intolerance scores were used to rank and prioritize the genes and not as cut-offs for selection.

After shortlisting variants according to the aforementioned criteria, we performed a literature review on the prioritized candidates and checked if coding variants in important oncogenes, tumor suppressor genes or autosomal dominant familial syndrome genes had been missed by the cut-offs of the pipeline. These variants were handled leniently with regard to conservation and deleteriousness cut-offs and were included in the further analysis.

### Candidate variant selection and validation

After filtering the variants based on the FCVPPv2, we visually inspected the WGS data for correctness using the Integrative Genomics Viewer (IGV) (Robinson, Thorvaldsdottir et al., 2017). The final selection was based on a thorough literature review. The selected variant of interest (*POT1 p.V29L*) was validated by Sanger sequencing of DNA samples of all available family members using specific primers for polymerase chain reaction amplification designed with Primer3 (http://bioinfo.ut.ee/primer3-0.4.0/). Primer details are available on request. Sequencing was performed on a 3500 Dx Genetic Analyzer (Life Technologies, CA, USA) using ABI PRISM 3.1 Big Dye terminator chemistry according to the manufacturer’s instructions. The electrophoretic profiles were analyzed manually. Segregation of the variant with the disease was confirmed.

### Further *in silico* studies

SNAP2 (Hecht, Bromberg et al., 2015), a neural network-based classifier, was accessed via PredictProtein (Yachdav, Kloppmann et al., 2014) to generate a heat map representation of independent substitutions for each position of the protein, based on its tolerance to amino acid substitution. The effect of the p.V29L mutation on the stability of the POT1: DNA interaction was assessed using the mutation Cutoff Scanning Matrix (mCSM) tool (Pires, Ascher et al., 2014).

### Protein alignment and structural modeling

Multiple sequence alignments were generated for homologous POT1 sequences to evaluate conservation using T-Coffee (Notredame, Higgins et al., 2000). Alignments for POT1 were generated using the following sequences: NP_056265.2, XP_519345.2, NP_001127526.1, XP_009001386.1, XP_006149256.1, NP_598692.1, XP_002712135.2, XP_010802750.1, XP_005628494.1, XP_001501458.4, XP_006910616.1, XP_010585693.1, XP_004478311.1, XP_007504310.1, XP_001508179.2, NP_996875.1, and NP_001084422.1. Alignments were visualized and formatted manually.

### Cell culture

HEK293T cells (RRID: CVCL_0063) were a gift from Andreas Trump (DKFZ, Heidelberg). The cells were maintained in DMEM high glucose supplemented with 10% fetal bovine serum (Gibco), penicillin (50 U/ml, Life Technology), and streptomycin (50 µg/ml, Life Technology). The cells have been authenticated using SNP or STR profiling within the last 3 years and all experiments were performed with mycoplasma-free cells.

### Selection and expansion of stable POT1^WT^ and POT1^V29L^ clones

pLPC myc hPOT1 was a gift from Titia de Lange (Addgene plasmid # 12387; http://n2t.net/addgene:12387; RRID:Addgene_12387; (Loayza & De Lange, 2003)). *POT1*^*wt*^ and *POT1*^*V29L*^ plasmids were transfected into HEK293T cells at 70–80% confluence using Lipofectamine 2000 (Thermo Fisher Scientific). At 48 hours post-transfection, cells were selected by growth with 1 μg/ml of puromycin for 14 days. The medium was changed every 2-3 days. Surviving colonies were selected and used for further experiments.

### Measurement of relative telomere length

Telomere length was measured on DNA extracted from HEK293T *POT*^*WT*^ and *POT1*^*V29L*^ cells after 40 passages using real-time PCR as described earlier by others and in our lab (Hosen, Rachakonda et al., 2015). Telomere and albumin primer sequences 5′ to 3′ were: ACACTAAGGTTTGGGTTTGGGTTTGGGTTTGGGTTAGTGT (Telg), TGTTAGGTATCCCTATCCCTATCCCTATCCCTATCCCTAACA (Telc), CGGCGGCGGGCGGCGCGGGCTGGGCGGCCATGCTTTTCAGCTCTGCAAGTC (Albugcr2) and GCCCGGCCCGCCGCGCCCGTCCCGCCGAGCATTAAGCTCTTTGGCAACGTAGGTTTC (Albdgcr2). Telomere/single-copy gene (T/S) values were calculated by 2^−ΔCt^ and relative T/S values (i.e., RTL values) were generated by dividing sample T/S values with the T/S value of reference DNA sample (genomic DNA pooled from 10 healthy individuals). All the experiments were done in triplicates and repeated twice.

### Western Blot

Protein lysates were prepared and quantified using the BCA protein assay kit (Pierce, Darmstadt, Germany). 20 μg of the proteins were then blotted onto 0.2 μM nitrocellulose membranes and blocked with 5% milk. Membranes were incubated overnight at 4°C with the target Anti-Myc tag antibody [9E10] - ChIP Grade (ab32). Immune complexes were detected with the corresponding HRP-conjugated secondary antibody (Anti-rabbit IgG, HRP-linked Antibody, cell signaling, 7074). The loading quantity control was incubated with the Anti-beta-Actin antibody [AC-15] (HRP) (ab49900) overnight at 4°C. Blots were developed by using ECL Western blot substrate (EMD Millipore, Darmstadt, Germany).

### Chromatin Immunoprecipitation (ChIP) assay and telomere dot-blots

The ChIP assay was performed following the protocol by Liu F. et al. with minor modifications (Liu, Feng et al., 2017). Briefly, the cells were cultured in 15 cm^2^ dishes with 70% confluence and fixed for 10 min at 25°C with a working solution of 1% (v/v) formaldehyde on a shaking platform. The cross-linking reaction was quenched by adding glycine to a final concentration of 0.125 M. ChIP was performed using the Chromatin immunoprecipitation assay kit (EMD Millipore, #17-295) following the manufacturer’s instructions using 5 µg of a mouse monoclonal to Myc tag – ChIP Grade antibody (Anti-Myc tag antibody [9E10] - ChIP Grade ab32). The lysates were sonicated with 5 × 5 min with 5s on/off intervals (Bioruptor, Diagenode) to get the DNA lengths between 200 and 1000 bp. The immunoprecipitated DNA was purified with the iPure kit (Diagenode, c03010014). Purified DNA was slot blotted onto a Hybond N+ membrane with the help of a dot-blot apparatus (Bio-Rad, 170-6545) and subsequently hybridized with a biotin-labeled (TTAGGG)_3_ probe synthesized by Sigma. The North2South^®^ Chemiluminescent Hybridization and Detection Kit (Thermo Fisher: 17097) was used to detect the biotin signal with the help of a CCD camera. Signals were then quantified by Image J, and the fold of enrichment was calculated. The amount of telomeric DNA after ChIP was normalized to the total input telomeric DNA.

### Electrophoretic mobility shift assay (EMSA)

The gel shift assay of *POT1*^*wt*^ *and POT1*^*V29L*^ was performed as described previously (Ramsay et al., 2013). In brief, 20 µl reaction was prepared in EMSA buffer (25 mM HEPES-NaOH (pH 7.5), 100 mM NaCl, 1 mM EDTA and 5% glycerol) supplemented with 1 µg of poly(dI-dC) and around 30-40 ng of γP32 labeled ds telomere Probe (GGTTAGGGTTAGGGTTAGGG) per reaction. The reactions were incubated for 30 min. at 25°C. POT1^wt^ and POT1^V29L^ were immunoprecipitated from HEK293T cells, ectopically expressing respective *POT1* proteins, by lysing them in 20 mM Tris pH 7.5, 40 mM NaCl, 2 mM MgCl_2_, 0.5% NP40, 50U/ml Benzonase, supplemented with protease and phosphatase inhibitors. After 15 min. of incubation on ice, the NaCl concentration was adjusted to 450 mM and the incubation was continued for another 15 minutes. Lysates were clarified by centrifugation (13200 rpm, 20 min, 4°C) and 1.0 mg of total protein was used per immunoprecipitation in IP buffer (25 mM Tris-Cl (pH 7.5), 150 mM NaCl, 1.5 mM DTT, 10% glycerol, 0.5% NP40) supplemented with protease and phosphatase inhibitors. Endogenous proteins were captured onto protein G-magnetic beads (NEB; #S1430S), washed extensively in IP buffer and used for *POT1*^*wt*^*and POT1*^*V29L*^ source. After gel shift incubation, the reaction contents were loaded onto a pre-electrophoresed 5% acrylamide/bis (37.5:1) gel in 0.5xTBE and run at 100V at 25°C. The gels were dried and analyzed by autoradiography. The labeled probe consensus alone served as a negative control of the EMSA.

## Acknowledgments

The authors thank the members of the families for participating in this study, Genomics and Proteomics Core Facility (GPCF) of the German Cancer Research Center (DKFZ) for providing excellent library preparation and sequencing services and the Omics IT and Data Management Core Facility (ODCF) of the DKFZ for the whole genome sequencing data management. KH was supported by the EU Horizon 2020 program grant No. 856620.

## Author contributions

K.H., A.F. and O.R.B. conceived and designed the study. E.B. provided the NMTC family samples. N.P. ran the WGS pipeline. A.K., A.S., B.M., N.P., and O.R.B., analyzed the data. A.S., B.M., D.S. and V.K. performed the experiments. A.S. and O.R.B. wrote the first draft of the manuscript. All authors read, commented on and approved the manuscript.

## Conflict of interest

The authors declare no conflict of interest

## The Paper Explained

### Problem

Familial non-medullary thyroid cancer (FNMTC) accounts for 3–7% of all thyroid cancers with the transmission pattern being an autosomal dominant mode of inheritance. The risk of developing cancer for first-degree relatives is increased 3-5-fold compared to the general population and the genetic alterations responsible for this disease are hardly known. Several low-penetrance predisposing loci have been identified using GWAS studies in recent years, but no high-penetrance mutations have been confirmed so far.

### Results

We identified a novel germline variant in the *POT1* gene, a component of the shelterin complex. Functional *in vitro* studies showed that the mutation weakens the binding of POT1 to telomere DNA, leading to abnormal telomere elongation. This is the first *POT1* mutation identified in a family with a predominance of thyroid cancers, thereby expanding the spectrum of cancers associated with mutations in the shelterin complex.

### Impact

Identification of *POT1* mutations associated with familial non-medullary thyroid cancers is important, as it provides the possibility of early diagnosis and screening of individuals at risk of developing NMTCs and enables the development of personalized therapy.

## For More Information

Supplementary tables

Supplementary table 1: Short-listed variants with scores

Supplementary table 2: Known *POT1* germline variants

